# Comparing Brain-Score and ImageNet performance with responses to the scintillating grid illusion

**DOI:** 10.1101/2025.06.18.660291

**Authors:** Martin Kent Kraus, Lucy Verkerk, Sander Keemink

**Affiliations:** Department of Machine Learning and Neural Computing, Donders Institute for Brain, Cognition and Behaviour, Radboud University, Nijmegen, The Netherlands

**Keywords:** Perceptual illusions, Brain-Score, scintillating grid illusion, representational similarity

## Abstract

Perceptual illusions are widely used to study brain processing, and are essential for elucidating underlying function. Successful brain models should then also be able to reproduce these illusions. Some of the most successful models for vision are several variants of Deep Neural Networks (DNNs). These models can classify images with human-level accuracy, and many behavioral and activation measurements correlate well with humans and animals. For several networks it was also shown that they can reproduce some human illusions. However, this was typically done for a limited number of networks. In addition, it remains unclear whether the presence of illusions is linked to either how accurate or brain-like the DNNs are. Here, we consider the scintillating grid illusion, to which two DNNs have been shown to respond as if they are impacted by the illusion. We develop a measure for measuring Illusion Strength based on model activation correlations, which takes into account the difference in Illusion Strength between illusion and control images. We then compare the Illusion Strength to both model performance (top-1 ImageNet), and how well the model explains brain activity (Brain-score). We show that the illusion was measurable in a wide variety of networks (41 out of 51). However, we do not find a strong correlation between Illusion Strength and Brain-Score, nor performance. Some models have strong illusion scores but not Brain-Score, or vice-versa, but no model does both well. Finally, this differs strongly between model types, particularly between convolutional and transformer-based architectures, with transformers having low illusion scores. Overall, our work shows that Illusion Strength measures an important metric to consider for assessing brain models, and that some models could still be missing out on some processing important for brain functioning.

## Introduction

Image classification, the task of correctly labeling what is present in an image, has long been a staple of Artificial Intelligence (AI) research. Its applications are diverse, from self-driving cars to helping identify cancer (Wildi et al., 2022; Sharma et al., 2021).In many of these tasks, Deep Neural Networks (DNN’s) specializing in image classification match or outperform humans (Dodge & Karam, 2017; Sharma et al., 2021). In addition, although the networks are not trained to do so, many of these deep neural networks also have a strong match between their neural activations, and those found in brain measurements (Schrimpf et al., 2018). Moreover, image labeling behavior can also be quite similar to humans both in terms of successes and in failures, in particular under time pressure (Elsayed et al., 2018). This suggests that these networks are not just good AI tools, but also that the representations learned by the model for image processing may be similar to those of the brain. Strengthening this view, even several perceptual illusions have been shown to be measurable in deep neural networks (Watanabe et al., 2018; Ward, 2019; Ngo et al., 2023; Zhang & Yoshida, 2024). Since the presence of visual illusions could indicate specific statistical assumptions about the world (Gregory, 1980; Tyler, 2022; Palmer & Rock, 1994), a good model of the brain should then not only be able to predict brain responses accurately, but also share the assumptions which lead to the presence of illusions. However, most previous illusion studies only considered a limited number of networks, and generally did not correlate illusion measurement with brain predictivity nor model performance. In this paper we aim to study this in more depth for a single illusion: the scintillating grid illusion (Schrauf et al., 1997) (Fig. 1).

**Figure 1.**
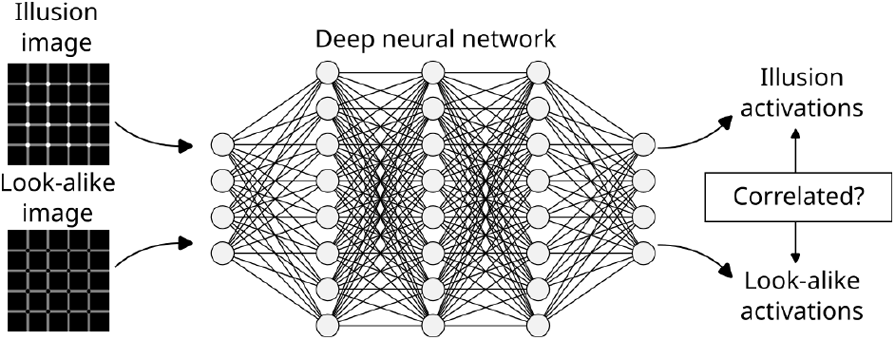
Deep neural networks trained for image classification correlate well in their labeling behavior and neural activations with brain and behavioral measurements. Visual illusions can sometimes also be measured, in this case by comparing layer activations to illusion and look-alike images. If the responses are similar in later layers, this suggests that in the processing of this image the network also ‘perceives’ the same illusion.

Many illusions have been studied in DNNs. These include the snake illusion (Watanabe et al., 2018), the tilt illusion (Benjamin et al., 2019), various contrast illusions in networks trained for de-noising and de-blurring (Gomez-Villa et al., 2020), the Vasarely illusion (Ngo et al., 2023), the MüllerLyer illusion (Ward, 2019; Zhang & Yoshida, 2024)and the Brightness Contrast Illusion in Diffusion models (Gomez-Villa et al., 2024).A study of illusions in a Visual Language Models has shown their propensity to identify illusions even where there are none (Ullman, 2024). A common approach in these studies is to measure the activations in response to an illusory image and then interpret these activations. However, many illusions involve both low-level and high-level processing, and generally require the subject to be able to give highlevel answers such as ‘what is the color of this object?’ or ‘what orientation do you see for this specific line?’. This is not always possible for standard deep neural networks. An alternative is to use representational similarity, in which illusion measurements can instead be achieved through comparing how neural activations in DNNs are correlated between illusion images and look-alikes (Fig. 1). The advantage of this approach is that it is both more model and illusion agnostic. This approach has recently been successfully applied for several illusions (Ward, 2019; Sun & Dekel, 2021; Zhang & Yoshida, 2024), which produced mixed results in finding the presence of illusory responses in networks.

We will consider the scintillating grid illusion as it was recently been shown to exist for several DNNs. In addition, responses from human participants were also obtained using the same methodology (Sun & Dekel, 2021).In the scintillating grid illusion, black circles appear at the center of the white disks in our peripheral vision as our gaze shifts. A set of images based on the Scintillating Grid were created, where the white disks gradually shifted towards black disks. A non-monotonic relationship was found in the representation of these images – images with black circles, close to what the illusion makes us see – were rated as more similar to images with white disks, than to images with gray circles. The central assumption of this methodology is that such non-monotonicity is a proxy for the presence of an illusion. Without an illusion, images with increasingly dissimilar pixels should result in responses with increasing dissimilarity to the original image. Models which experience the illusion should have more similar responses between the illusion and lookalike images (meant to ‘mimic’ the illusion), rather than with in-between images (Fig. 1).

Building on this previous work, we first defined a simple metric for Illusion Strength which takes into account both response similarity between illusion and control image responses (Fig. 1), and applied it to 51 different models, chosen to be representative of different model types. We investigated which kinds of networks show illusion-like representations, and found 41 out of 51 showed illusion-like representations. We next compared our metric with how well these models can predict both brain responses and image labeling behavior by using their Brain-Score (Schrimpf et al., 2018), and compared to their performance by considering the top 1 ImageNet accuracy (Deng et al., 2009). We originally hypothesized that better Brain-Scores would also lead to strong illusion scores, but we found no strong correlation between the strength of the illusory representations and Brain-Score or ImageNet accuracy, suggesting that Brain-Score and similar metrics could us illusion perception as an additional metric for how ‘brain-like’ models are. Finally, we show significant differences between how the Scintillating Grid is processed between different families of architectures.

## Methods

The hypothesis set out in Sun & Dekel (2021) is that for networks that experience the scintillating grid illusion similar to humans, we should see that the responses to images with black and near black disks look more similar to white disks, than those in between (making ‘black-disk’ images a ‘lookalike’). We here build on their analysis to test this property and compare it to controls. The code for running the experiments can be found at https://github.com/keemink-lab/sg illusion paper code main.

### Generating images

#### Illusion images

We retrieved the original images with white and black disks from Sun & Dekel (2021). The in-between images were created through blending together two images with the Python PIL library, as can be seen in Figure 2A. The illusion (white-disk image on the left) and the lookalike images (black-disk image on the right) were blended in steps of 5%, going from 100% illusion and 0% lookalike to 0% illusion and 100% lookalike, resulting in 21 luminance levels. In addition, to test for the illusion across different viewing conditions, the grids in these images varied in the number of lines, size of the disks and offset from the center. These variations are discussed in greater detail in the Sun and Dekel’s (Sun & Dekel, 2021) Image Stimuli section. In total, 375 variations were used, with 21 luminance levels for each variation.

**Figure 2.**
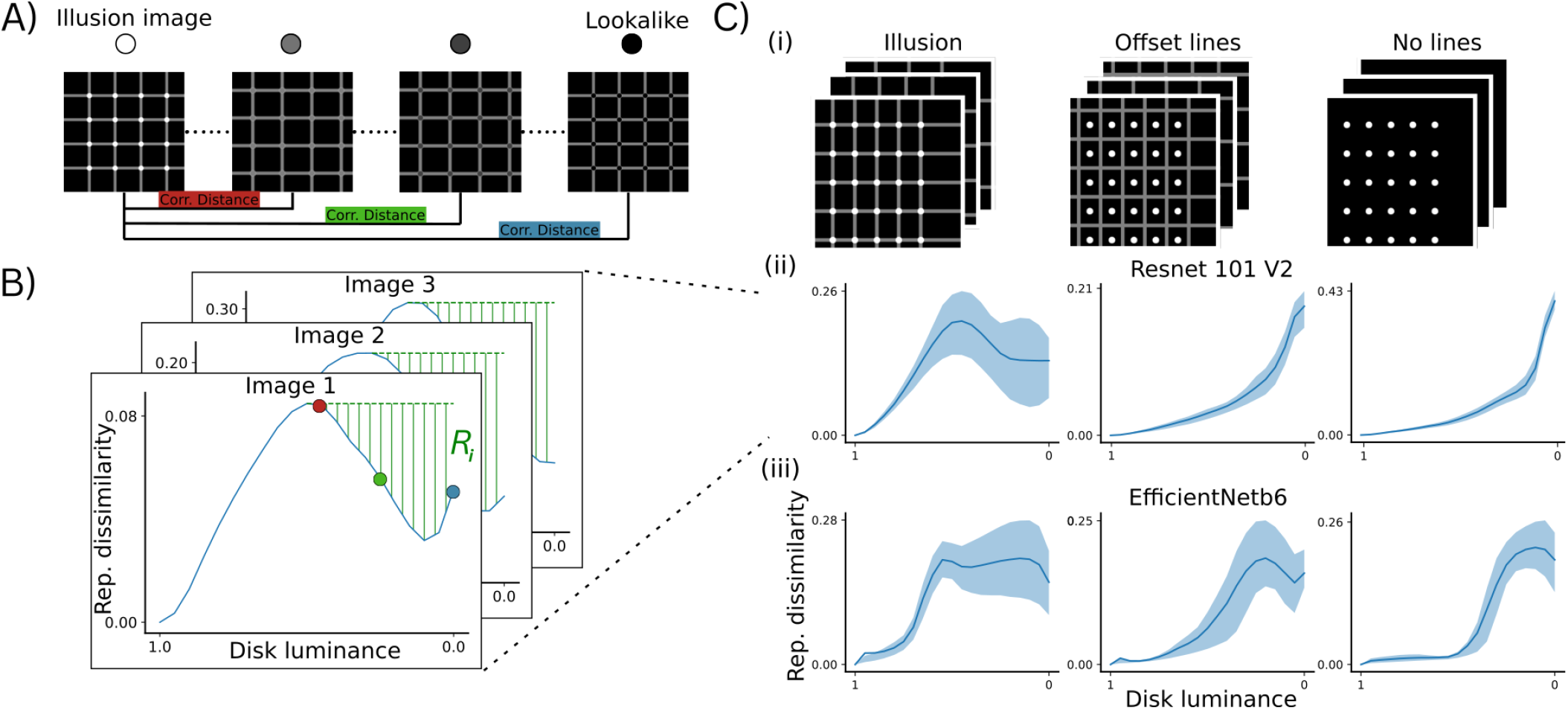
Measuring illusion response on scintillating grid and control images. A) The scintillating grid illusion makes small black disks appear inside the white intersections. To check for the presence of the illusion in a neural network, we create new images by transitioning the color of the circles from white to black. Each image is then presented to the network, with the activations in the penultimate layer being recorded. The Pearson correlation distance of the response to each image to the original image (with white disks) is then calculated. B) Determining the presence of an illusory response. We here plot the correlation dissimilarity between the illusion image (white disks) and other images. The red, green and blue dots correspond to correlation distances recovered between the images in panel A for Resnet-101. Since the only thing changing between the images is the color of the disks, for networks without an illusory response the expectation is a monotonically increasing curve (the image with the black dots is then the most different from the image with the white dots). If responses are non-monotonic, this suggests the model represents the illusion similarly to what the illusion makes us see. To check for illusion-like responses, we quantify the area between the highest Pearson correlation distance and the lowest luminance image, shown here as the dashed green area, as the deviation magnitude. C) Example average responses across different image sets. (i) From left to right, example illusion and control images. Controls are needed to check against black and white circles simply being represented similarly by the model. Each type of image has many permutations through offsets, disk sizes, and other changes (see main text). (ii) Average responses for Resnet 101’s penultimate layer across the images. The mean (curve) and inter-quartile (areas) are shown. Here, the network has a non-monotonic response to the illusion images, but a fully monotonic response to the controls — indicating an illusory response. (iii) Same as (i), but for EfficientNetB6. In this case there might be a weak illusory response to the illusion images, but also to the control images, as all curves are non-monotonic.

#### Control images

The same procedure as for the illusion images was used to generate two sets of control images, also taken from Sun and Dekel (Sun & Dekel, 2021) (Fig. 2Ci). These are the ‘No lines control’ where the lines of the grid are absent, and the ‘Offset lines control’ where the disks are moved from the intersections of the grid to the center of the four surrounding lines. These controls were chosen as humans see no illusory black disks on the white disks in these cases, and previously significant differences in network representations on them compared to the illusion images were found (Sun & Dekel, 2021). Each control was subjected to the same variations as the illusion image set, resulting in again 375 variations for each control with 21 luminance levels each.

### Measuring Illusion Strength

In order to analyze the internal representations of each network, activations were retrieved with the use of Pytorch hooks (Paszke et al., 2019) from the penultimate layer. This was done in order to replicate Sun and Dekel’s (Sun & Dekel, 2021) results, as they found non-monotonicity emerged gradually in the deeper layers of a network. We also reproduced this analysis for our networks, and found the same phenomena (shown only for some representative networks, Fig. 3). In addition, for convolutional networks, the penultimate layer has been shown to perform most similarly to humans when used as a model for various vision tasks (Jozwik et al., 2017; Kubilius et al., 2016) though this is not conclusive (Dekel, 2017).

**Figure 3.**
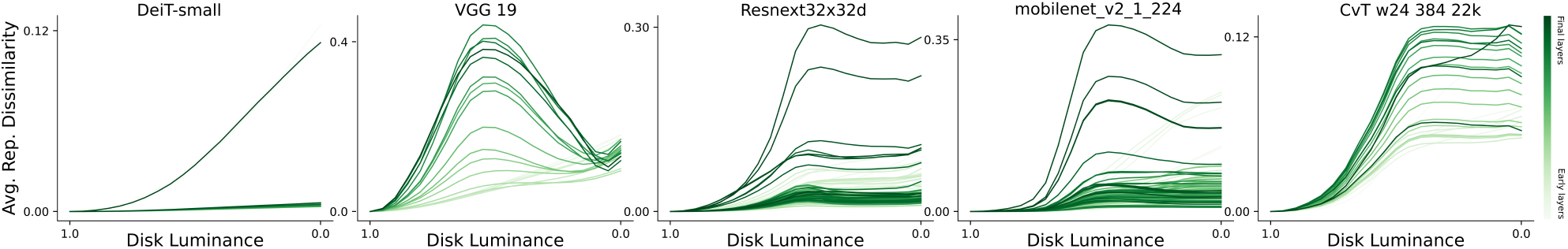
Comparing the average representational similarities across model layers, on illusion images. From left to right: representational similarity curves for the DeiT-small, VGG 19, Resnext32×32d, mobilnet v2 1-224, and CvT w24 384 22k networks. Lighter green color corresponds to earlier layers, and darker green color corresponds to later layers. Hooks were placed in the penultimate pooling or fully connected layers as well as earlier layers. These were chosen to be the convolutional layers for Resnext32×32d and mobilnet v2 1-224, a combination of convolutional and fully connected layers for VGG 19, Layernorm for CvT w24, and the projection layers for DeiT-small

Sun and Dekel originally used a normalized L1 distance, but as the normalization occurred per image set for each network it is non-trivial to compare the resulting values across different sets of images and networks. We instead quantify the dissimilarity between internal representations by the correlation distance between the responses to images. For each set of images, we take the activations of the penultimate layer on the reference image **x**_ref_, which can either be a control or contain an illusion, and compare to images with adjusted disk luminance **x**_comp_(*q*), where *q* is the disk luminance ranging from 1 to 0 (Fig. 2A). We compute the representational dissimilarity of the response between two images *C* as follows:

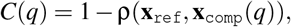

where ρ(**x**_1_, **x**_2_) measures the Pearson correlation. As such the representational dissimilarity *C*(*q*) is 0 if the images result in completely identical responses, or 1 if they are completely uncorrelated. Across images this results in a set of curves that show how the dissimilarity changes across luminances (Fig. 2B). When mapping out the dissimilarity curve across disk luminances, in the case of an illusory representation, we would expect to see a non-monotonic curve that first goes up and then comes down again.

After all pairs of illusion and lookalike images have been presented to the model, the deviation magnitudes for each network and each image set are computed (Fig. 2 B). We define the deviation magnitude as the total change from the highest point in the curve with all the following points as the presence of this non-monotonicity indicates a higher similarity between darker disks and white disks than those in-between. We compute the deviation from monotonicity, or deviation magnitude *R*_*i*_ as:

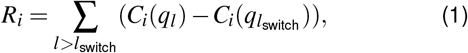

where *C*_*i*_(*q*_*l*_) is the correlation distance for image *i* and the *l*’th image with luminance 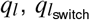 is the switching point at which images start to become less similar compared to the highest point in the curve (Fig. 2B). We choose this point because we expect that on images without illusions, increasingly different images should also be considered to be increasingly different by the model. If the relationship instead becomes non-monotonic - a decrease in correlation distance compared to previous-images then the model instead internally represents some (more dissimilar pixel-wise) images as being more similar conceptually. This measurement is computed per each illusion-lookalike pair, with the total deviation magnitude being defined as the the average 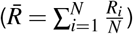 over all images in a set (either illusion, or one of the controls).

Following Sun and Dekel, the highest point was chosen as the reference point (rather than, say, the first deflection from monotonicity). This avoids cases where the black look-alike image has a very high dissimilarity, but where there is some other point on the curve before which corresponds to a non-monotonic reflection-point.

To conclude that a network has illusion-like representations, it is not enough to have a positive deviation magnitude on illusion images. Another requirement must be that nonmonotonicity is either significantly reduced or absent on control images. The need for this can be seen in Fig. 2 Ciii, where an example network has (on average) a stronger ‘illusion-like’ response on one of the controls than the illusion image. To take this into account we next define the ‘Illusion Strength’ measure for a given network as

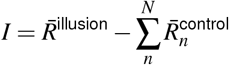

where the 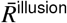 is the average deviation magnitude on the illusion set, *N* is the number of control image sets, and 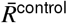 is the average deviation magnitude on the *n*’th control image set. We thus measure the illusion strength as the deviation magnitude on the illusion images, penalized by the deviation magnitudes on the control images. A positive illusion strength indicates that there was more deviation magnitude on the illusion images, than in total on the control images. A negative illusion strength indicates the opposite. We also considered a stricter measurement, where networks with any deviation magnitude on controls were classified as not having illusion-like representations.

### Models studied

Our study was conducted on 51 models. These were chosen from among those listed on Brain-Score.org (Brain-Score, 2018), in such a way as to cover a wide range of Brain-Score values, and to include commonly used architectures such as VGGs, ViTs and ResNets. We chose networks from the VGG (Simonyan & Zisserman, 2014), ResNet (He et al., 2016), WSL ResNext (Mahajan et al., 2018), DenseNet (Huang et al., 2017), MobileNet (Howard et al., 2017; Sandler et al., 2018), SqueezeNet (Iandola et al., 2016), DeiT (Touvron et al., 2021), Inception (Szegedy et al., 2016, 2017), EfficientNet (Tan & Le, 2019), ViT (Dosovitskiy, 2020) and CViT (Wu et al., 2021) families, as well as Alexnet (Krizhevsky et al., 2012).

### Brain and performance scores

Our study used two main scores to compare Illusion Strength with, namely the Brain-Score (Schrimpf et al., 2018) and the ImageNet top-1 score (Deng et al., 2009). The ImageNet top1 scores were taken from the websites hosting the model, such as Pytorch (Paszke et al., 2019), or from the respective papers of each model when this was not available. Brain-Scores were taken from brain-score.org (Brain-Score, 2018) on December 10th 2024^1^. For six models from the CViT family, scores were not available, and so were procured by submitting the models to the Brain-Score website and generating their scores. The Brain-Score was chosen as it represents an extensive summary of how well models predict both brain recording and behavioral data.

## Results

### Deviation magnitudes on illusion and control images

We first consider if networks experience ‘illusion-like’ responses to scintillating grid images, by comparing the original illusion image (with white disks) to images with progressively darker disks (Fig. 2A). We then measure the dissimilarity through correlating the responses to each image pair, and measure the ‘deviation magnitude’ as the average drop from the highest point the curve (Fig. 2B). This results in, for example, Resnet 101 having a significant deviation magnitude, as on illusion images it has a non-monotonic response (Fig. 2Cii, left). In contrast, networks that have a weak or no drop at all will have a low deviation magnitude, as for the EfficientNetb6 (Fig. 2Ciii, left). Overall, we find that 45 out of 51 networks show non-zero deviation magnitudes on illusion images (Fig. 4, x-axis).

**Figure 4.**
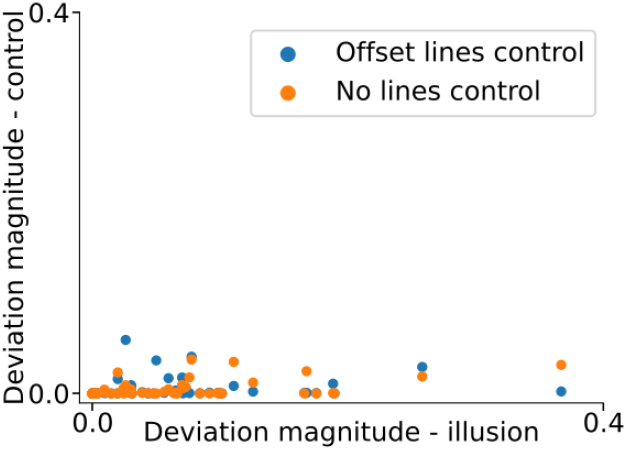
Comparing deviation magnitudes between illusion images and controls (‘offset lines control’ in blue, and ‘no lines ‘control’ in orange). See Fig. 2Ci for example images of the controls.

However, if a network shows this non-monotonicity also on images that do not elicit a illusory perception in humans, then the model just generally has such non-monotonic responses to the disks changing, not in the specific location we would expect the illusion to occur. To control for this, we additionally define two control set images, on which we apply the same procedure as for the illusion image (Fig. 2Ci).Here, we find strong differences between networks. Some networks, like Resnet 101, have purely monotonic dissimilarity curves for the controls, compared to a highly non-monotonic response to the illusion images (Fig. 2Cii). This indicates that these networks have uniquely illusory responses only to the illusion images. In contrast, other networks, like the EfficientNetb6, can have non-monotonic dissimilarity curves for the illusion images as well as the controls (Fig. 2Ciii).

When we compare these deviation magnitudes for all 51 networks tested, we see that most networks which have a deviation magnitude on the illusion images, also have them on the control images, to varying degrees (Fig. 4).While deviation magnitudes on the control images were usually much smaller, Resnext32×16d and Inception ResNet v2 had stronger deviation magnitudes on control images compared to the illusion images.

We then proceeded to determine whether the observed deviation magnitudes are significant. Because deviation magnitudes are non-negative, they are not normally distributed, and thus require a non-parametric test. We chose one-sided Mann-Whitney U tests, comparing whether the distribution of the 375 deviation magnitudes obtained on the illusion is larger than the deviation magnitudes obtained on a single control. We thus obtained two p-values, one for each control condition. Of these two, the higher p-value was chosen, and compared with a Bonferroni corrected significance level of 0.00049. As a result, the responses of 41 networks to the illusion were found to be significantly larger than the responses to both of the control sets. The significant models included VGG 19 and Resnet 101, which were also studied by Sun and Dekel (Sun & Dekel, 2021), by which we replicate their results.

### Investigating Illusion Strength

To take into account deviation magnitudes to both illusion and control images (Fig. 4), we next consider a measure for Illusion Strength which measures the deviation on illusion images, but penalizes it by the deviation magnitudes on control images (see Methods). An Illusion Strength of 0 indicates that even if the network did have some deviation magnitude on the illusion images, that this was also the case on the control images to the same degree.

When a more conservative measurement of Illusion Strength was used, which assigned 0 Illusion Strength to any network with any non-monotonicity on any control image, the number of networks with positive Illusion Strengths decreased to 3 out of 51, all of which were from the CViT family. However, given that many of these models only had very small deviation magnitudes on the controls, we considered this too strict as an illusion measurement.

After taking this into account, most networks - 41 out of 51 - were assigned a positive Illusion Strength score (Fig. 5). This suggests these models and by our measure at least have the scintillating grid illusion to some degree, but for them to be considered good brain models they must also match well behaviorally and in their responses. To test this we next turn to performance and brain predictivity measures.

**Figure 5.**
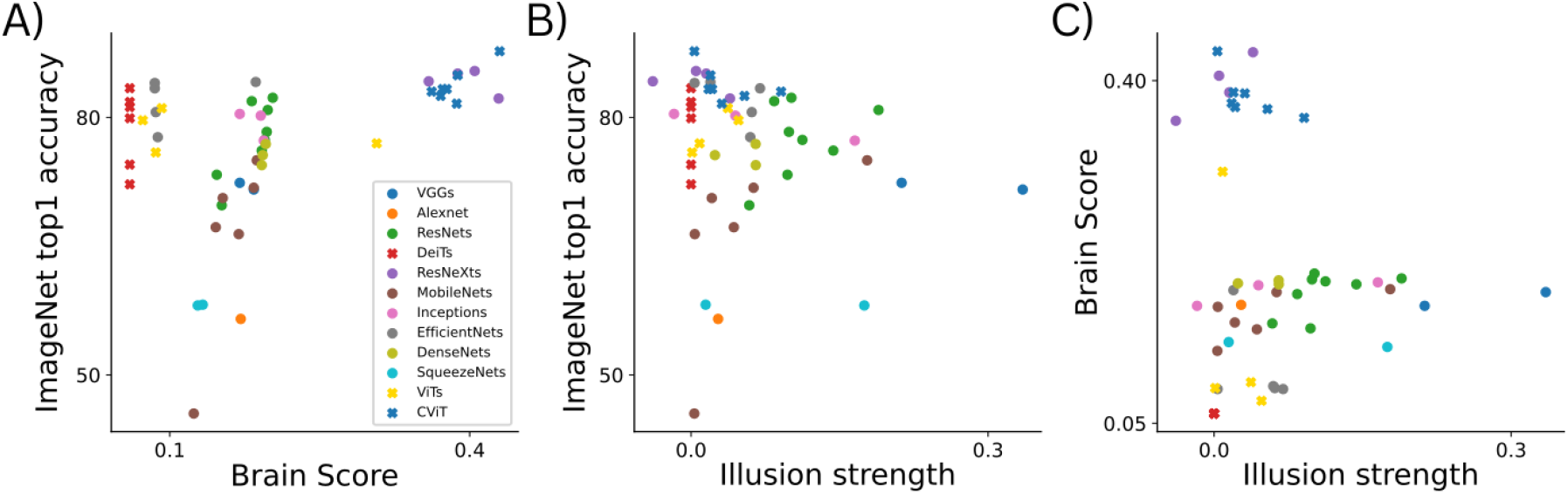
Comparing Brain-Score, ImageNet performance, and illusion strength. (A) Comparing the Brain-Score and ImageNet top1 accuracy scores for all models tested. Each color corresponds to a different model class (as indicated in the legend), and crosses correspond to transformer networks. (B) Same as panel A, but for comparing Illusion Strength to ImageNet performance. (C)Same as panel A, but for comparing Illusion Strength to Brain-score.

### Brain-Score and ImageNet performance

For a performance metric we will use the ImageNet top-1 Accuracy score, and for how well models predict brain and behavioral data we will use the Brain-Score measure. The Brain-Score is useful as it collates several behavioral and neural measurements, and is meant to assess a model for how good of a model for human vision it is. See Methods for a description of how we retrieved calculated or retrieved these scores.

We now first consider how Brain-Score and ImageNet performance relate to each other (Fig. 5A). While there is a Pearson correlation of *r* = 0.37 between Brain-Score and ImageNet top-1 scores in the considered networks, there does not seem to be a linear relationship, and three groups of models are visible. We observe that models with similar architectures cluster together. It is noteworthy that models from the CViT family and ResNeXts not only achieve good ImageNet accuracies as well as high Brain-Scores, but that they obtain the highest scores in both measurements.

### Comparing Illusion Strength with performance

We next consider if the presence of scintillating grid illusory responses relates to ImageNet Performance (Fig. 5B).We measure a negative correlation between the performance of the models and their Illusion Strength (*r* = −0.15). However, this is nonconclusive as we also observe a strong heteroscedasticity. We observe that the variance is not constant throughout, and increases particularly for models with low performance and high Illusion Strength. Therefore, the Pearson correlation reported above over-estimates the strength of the relationship. However, it is clear that the top-performing models classes (CViTs and ResNeXts) both have very low Illusion Strength.

### Comparing Illusion Strength with brain-score

Finally, we look at how well Illusion Strength correlates with brain predictivity through Brain-Score (Fig. 5C). When plotted against each-other, this results in a complex distribution, seemingly divided into two clusters, with no strong positive or negative correlation (*r* = −0.06) between Illusion Strength and BrainScore. It is however of interest that only models which score relatively low on the Brain-Score exhibited strong Illusion-Like responses. When CViTs and ResNeXts are omitted, we instead observe a moderate relationship of *r* = 0.39.

We generally saw that models from the same family of architectures clustered together. There were four families of architectures where all models showed illusion-like responses: VGGs, ResNets, SqueezeNets and CViTs. This seems to be a particularly robust result for ResNets, where eight models were measured. On the other hand, Transformer based architectures without convolution showed much smaller illusion-like responses. Of the 10 non-convolutional Transformer models studied, 7 did not have strong illusion-like responses. The DeiT architectural family, consisting of six networks, was the only one where no networks had any illusion-like responses.

### Network responses across luminances

We additionally investigated how different networks represented the Scintillating Grid across all disk luminances using Representational Dissimilarity Matrices (RDMs). Doing this analysis allows us to investigate in more detail how the different networks might end up with an illusory response, and if the non-monotonicity is unique to the white and close-to-black disks, or if there are other patterns. We here plot the RDMs for a few representative patterns from the full dataset.

Representational Dissimilarity Matrices revealed five patterns in the internal representations of the networks on the Scintillating Grid images (Fig. 6). In networks with no Illusion-Strength, such as DeiTs (Fig. 6 first panel), increasingly pixelwise distant images were represented as increasingly dissimilar, resulting in black diagonal in the middle. For VGGs and ResNets, which have strong illusory responses, images with minor color differences were represented similarly, and images with gray disks were equally distant to both the illusion and the lookalike images. However, images with white and black disks look very similar to each other, resulting in a cross-like shape (Fig. 6 second panel). For others, such as EfficienetNets and WSL ResNexts (Fig. 6 third panel), images with gray disks were represented much closer to the lookalike than the illusion image, resulting in a dual square shape.

**Figure 6.**
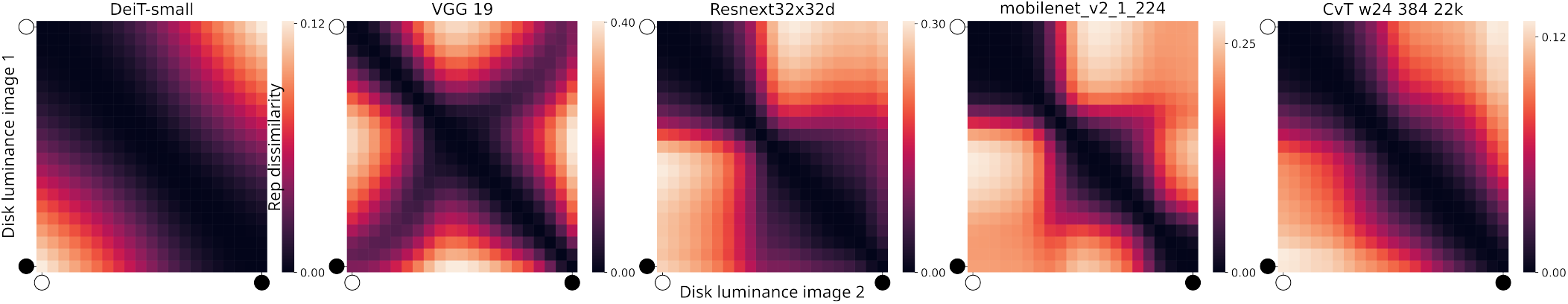
Comparing representational dissimilarity between all disk luminances, through Representational Dissimilarity Matrices (RDMs) on illusion images. The brighter the color, the more dissimilar the representation. Each row and column represents an image with a different disk luminance, such that the diagonal compares images with themselves (and so have a dissimilarity of 0). The top row corresponds to the curves drawn in Fig. 2B, as it compares images with white disks to all other luminances. Shown are RDMs for the DeiT-small, VGG 19, Resnext32×32d, mobilnet v2 1-224, and CvT w24 384 22k networks.

In MobileNets white, grey and black disks form distinct categories (Fig. 6 fourth panel) resulting in several square subsections. Finally, in the example CvT there is a waist like shape, indicating at least two categories of brighter and darker images — but there is no clear illusory response (Fig. 6fifth panel).

The above changes in representational dissimilarity between images with different reflect changes in how those images are being represented in the penultimate layer, and how it changes with disk luminance. For models in which there is a simple identity-like structure the mapping is monotonic: as the images become more and more dissimilar in pixel-space, so too do the network responses (as is the case for he DeiT-small network). For models in which the the RDMs show more detailed structures, the representations change more nonlinearly and non-monotonically across disk luminances (as for the other models shown). For the cross-like patterns of models such as VGG 19 the interpretation is that there is an illusion present similar to what is seen by humans (as for the VGG 19 example), but the other RDM patterns show that different representations are also possible even if not consistent with the illusory percept expected from human psychophysics. We also found that the representations (as one would expect) act much more monotonically in earlier layers than later layers (Fig. 3), and it would be of interest for later work to combine these analyses to figure out how illusory representations can manifest across layers.

## Discussion

Deep neural networks that are considered good models for human vision should also reproduce human visual illusions. While for some networks and some illusions this was established before (e.g. Zhang & Yoshida (2024); Benjamin et al. (2019); Ngo et al. (2023)), it was not clear how well this translates across a larger set of networks, nor how it correlates with brain predictivity. In this paper we studied responses to the scintillating grid illusion, and found that many networks indeed have an illusion-like response. Compared to previous work, we analyzed many more models than previous studies, and also directly compare brain predictivity with the illusion score. We did not find a strong correlation between brain predictivity (Brain-Score) or performance (ImageNet) and the presence of the illusion. In fact, we found a group of models (ResNeXts and CViTs) that perform very well on Brain-Score and ImageNet, but do not have illusion-like responses. Secondly, we had a group of models that had a strong illusion score (in particular VGGs), but performed relatively low on Brain-Score. This suggests that there are still models to be developed that perform well on reproducing the scintillating grid illusion, while also predicting brain responses well.

Our results rely on a measure of Illusion Strength inspired by Sun & Dekel (2021), but in addition taking into account responses to control images. The measure makes the implicit assumption that the illusion can be measured in networks by comparing the responses between illusions and look-alike im-ages. The differences in models are intriguing, in particular that transformers tend to have a smaller illusion scores than other networks. However, there are also many classic convolutional networks that also have low Illusion Strength, so the illusion-like responses are not just an artifact of convolution operations.

It is somewhat surprising that the scintillating grid illusion can be detected in deep neural networks, as for humans the illusion strongly depends on eye movements (Ninio & Stevens, 2000; Schrauf et al., 1997), which none of the studied networks explicitly model. By considering many permutations of the illusion images (e.g. in grid size) we did make sure that the responses were stable for different viewing conditions, and the fact that we (and Sun & Dekel (2021)) found strong differences between illusion image and controls further strengthens the validity of our result. It might be that the illusion is present for both humans and these networks through completely different mechanisms, but for similar underlying computational reasons.

The way we measured the illusion in this paper, and how it is measured in humans, is also significantly different. The assumption of non-monotonicity underpins the calculation of deviation magnitude and thus Illusion Strength. There has been some work measuring perceived illusion strength on the Scintillating Grid which indicates that non-monotonic responses may be the case (Schrauf et al., 1997; Sun & Dekel, 2021). However, perceived illusion strength in humans is not the same as the representational dissimilarity used here, and direct comparisons would assume similarities that may not be there. Therefore, a human study should be conducted to determine the validity of the assumption of non-monotonicity in the representations of illusions (for example by replicating our network experiment with RDMs, and asking participants to assess image similarity). It could then be determined which kinds of illusion RDMs are more brain-like. In addition, it would be of interest to determine which part of the visual system gives rise to the illusion, particularly since lateral-inhibition in the early visual system does not seem to be the mechanism behind this illusion (Geier et al., 2008; Sun & Dekel, 2021). Knowledge of the mechanisms of the illusion in the brain would allow for a more principled selection of the layers studied in DNN’s. Models which are truly similar to the brain should not only show similar illusory representations in general, but should also show neural correlates in comparable layers.

The field of psychophysics has many theories about the possible causes of perceiving illusions, as well as many examples where the illusion image is slightly manipulated so as to remove the illusion or alter its strength. For the Scintillating Grid, the effects of disk size, shape, orientation (Qian et al., 2009; Schrauf et al., 1997; Matsuno & Sato, 2019) as well as other features (Levine & McAnany, 2008; Sun, 2019) on Illusion Strength have been studied. Each of these studies provides an additional point on which comparison between DNNs and humans can be made, and the underlying assumptions of DNNs examined. Moreover, it would allow us to select models which match human behavior more closely. Another recent development is the use of language vision models (Ullman, 2024), where one can directly show the model an image and ask questions about it. While promising, this can lead to conflicting results (e.g. seeing illusions where there are none), and current language models are still hard to interpret directly — making the representational similarity approach a useful possible addition.

The fact that Illusion Strength was measurable in many networks, but that no single network both had a high BrainScore as well as a high Illusion Strength could suggest different things. It may suggest that while we find the scintillating grid illusion in these networks, the mechanism and reason for their existence is different for humans and the networks. It may also be that the networks are missing out on some core processing that the brain is doing, and we could still get high Brain-Score measures plus a strong Illusion Strength by making the models more ‘brain-like’. It could also indicate that we need a different way to assess illusion strength. Finally, it could also indicate that the Brain-Score measure is simply incomplete, and psychophysical measures (like our Illusion Strength) should be taken into account to further differentiate models for how well they model the brain. For this, future work should include many more illusions which can be measured across model architectures, and eventually be integrated in the brain-predictivity benchmarks. It could then also be necessary to define new ways to measure the illusions in humans, which allow for a fairer comparison between model and brain representations.

## Footnotes

1 Web archive link

